# Enformation Theory: A Blueprint for Evaluating Deep Learning Models in Genomics

**DOI:** 10.1101/2024.09.03.611127

**Authors:** Eyes S. Robson, Nilah M. Ioannidis

## Abstract

The nascent field of genomic AI is rapidly expanding with new models, benchmarks, and findings. As the field diversifies, there is an increased need for a common set of measurement tools and perspectives to standardize model evaluation. Here, we present a statistically grounded framework for performance evaluation, visualization, and interpretation using the prominent sequence-based deep learning models Enformer and Borzoi as case studies. The Enformer model has been used for applications ranging from understanding regulatory mechanisms to variant effect prediction, but what makes it better or worse than precedent models? Does its follow-up, Borzoi, offer improved performance and more informative embeddings as well as finer resolution? Our goal is to propose a general blueprint for answering such questions and evaluating new models. We start by contrasting the few-shot performance of Enformer and Borzoi to precedent models on the `GUAN`in`E` benchmark, which emphasizes complex genome interpretation tasks. We then examine Enformer and Borzoi intermediate embeddings in model-subjective principal component space, where we identify limiting aspects that affect model generalization. Finally, we present an interpretable decomposition of Enformer and Borzoi, which allows for global model interpretability and partial ‘backtracking’ to explanatory causal features. Through this case study, we illustrate a new protocol, Enformation Theory, for analyzing and interpreting deep learning models in genomics.

## Introduction

AI models are rapidly taking a leading role in genome interpretation and comprehension. Sequence-based genomic deep learning models [61, 26, 25, 2, 1, 28], for example, have the potential to predict the consequences of genome sequence variation and enable the design of new genomic sequences with desired properties. However, our understanding of these sequence-based deep learning models is in its infancy, and the field needs a standardized protocol for introducing new models and systematically analyzing and interpreting their results. Here, we provide a case study for analyzing new models using our proposed blueprint, Enformation Theory. It consists of three distinct, but interrelated dimensions of analysis:

- **Performance:** Evaluate model performance on a variety of tasks using an existing or novel public benchmark compared to previous models.
- **Visualization:** Explore model embeddings and concomitant datasets in low-dimensional space to validate assumptions and generate hypotheses.
- **Interpretation:** Interpret a model globally to elucidate *what* the model has learned and *why* it succeeds or fails [31].

While published models typically undergo some subset of these types of analyses, it is important to have a standardized set of thorough evaluations, benchmarking tasks, visualizations, and interpretability analyses that are applied to all newly published models to directly compare their understanding of genomic sequences.

This study focuses on Enformer [1] and Borzoi [28], applying the above protocol with a proposed standard set of analyses. For performance, we make use of the genomic AI benchmark `GUAN` in`E` [43]. We visualize sequences in low-dimensional space, highlight the appropriateness of benchmarks like `GUAN` in`E` , and identify interpretable variables (IVs) that underlie model performance through the use of our proposed interpretability analysis. `GUAN` in`E` uniquely allows us to examine the performance of Enformer and its follow-up Borzoi on a collection of **large-scale** human and eukaryotic genomic tasks designed for robustness and controlled evaluation.

For our global interpretability analysis [31], rather than relying on aggregated feature attribution approaches that operate on a per-example basis for *local* explainability [29, 31], we opt for a sophisticated decomposition of variance explained first proposed by Dinga et al. [12]. This theory-backed [17] approach allows for the use of exogenous, interpretable variables (IVs) in a functional decomposition, which we refer to as interpretable variable decomposition (IVD). IVD dramatically lowers the computational and technical barrier to model investigation and auditing, while enabling the use of statistical significance testing for increased methodological rigor.

### Background

The standard approach to genomic AI involves the intersection of traditional bioinformatics with approaches from computer vision, particularly convolutional neural networks (CNNs). Supervised convolutional models attempt to capture sequence motifs, similar to Position Weight Matrices (PWMs) [3, 61, 2], and higher-order interactions between them. To expand the input sequence context, some supervised models also employ dilated convolutions, typically strided over an extended input [26, 25]. Recent, yet still relatively underexplored, approaches incorporate self-supervised autoencoders pretrained on tokenized sequences from (multi)organismal corpora [24, 60, 9, 43].

Regardless of methodology, scoring and validating genomic AI models is a recurring issue [30, 9, 43], and it requires careful consideration of data leakage, such as average tissue methylation [13], across tracks [46] or organisms [1]. Here we apply our proposed protocol for model evaluation to Enformer [1] and Borzoi [28], utilizing the `GUAN` in`E` benchmark [43] and a functional decomposition of variance approach [12], each introduced briefly below.

#### Enformer

Enformer [1] is a powerful^1^ genomic AI model that processes 196, 608 base pairs (bp) of input sequence context. It uses compressive convolutional blocks whose embeddings are fed into a transformer [55] at 128bp resolution. Enformer was trained in a supervised fashion on 5,313 human and 1,643 mouse experimental tracks, including cap analysis of gene expression (CAGE) data, using only reference genomes for each organism. Not including the output head for the mouse, the Enformer totals 246.1M parameters, of which the human output head is ∼ 6.6%.

#### Borzoi

Borzoi [28] is a larger-context and higher-resolution follow-up to Enformer. Borzoi passes a single large context size of 524, 288 bp through convolutional blocks into a transformer architecture, whose output is upsampled into 32bp segments in the manner of U-Net [44]. This approach allows for additional supervision using RNA-seq data – an information-rich source of gene expression data previously difficult to model with genomic AI [62]. To compensate for its large, bottleneck-inducing input size, Borzoi is a leaner 185.9M parameters, of which its human output layer makes up nearly 7.9%.

#### GUANinE benchmark

Although AI benchmarks are not yet comprehensive nor widely adopted in genomics, they are the canonical approach for model comparison and evaluation in other AI fields [11, 56, 57, 40, 5]. They offer a systematic means to evaluate and describe a model’s successes and failures and have seen a gradual yet growing rate of creation and adoption in genomic AI as well [9, 43, 15, 30, 63, 34].

`GUAN` in`E` [43] is a large-scale AI benchmark designed for genomic sequence-to-function deep learning models. Compared to smaller-scale benchmarks, where single-task, highly specialized baselines can be difficult to properly train due to small sample sizes [27, 15, 30], `GUAN` in`E` allows approaches based on transfer-learning using multiask models like Enformer and Borzoi to be directly compared with deep, specialized genomic machine learning algorithms trained specifically for the benchmarking tasks. As a result, `GUAN` in`E` combines the best features of large-scale benchmarks in computer vision [11] with multitask benchmarks in natural language processing [56, 57, 52].

`GUAN` in`E` v1.0 includes region-level analyses such as genome-wide DNase hypersensitivity estimation, sequence conservation annotation, and large-scale synthetic gene expression reporter assays [43]. This makes `GUAN` in`E` an ideal choice for both evaluating model fit and articulating model competencies. The primary limitation of v1.0 is the lack of variant interpretation tasks or tasks specific to particular cell types. `GUAN` in`E`’s three distinct task groups vary in source organism: functional elements (human), conservation (human), and gene expression (yeast). The functional elements tasks include dnase-propensity and ccre-propensity, which ask the model to estimate the cell-type-agnostic^2^ likelihood that a sequence is accessible (DNase hypersensitive) and the cell-type-agnostic likelihood that an accessible sequence exhibits H3K4Me3, H3K27ac, or CTCF epigenetic features (indicative of promoters, enhancers, and cohesin loops, respectively). The cons30 and cons100 tasks ask a model to annotate the region-level (mean) PhyloP [38] score for 512bp, while the gene expression task is borrowed from Vaishnav et al. [54]. It is worth acknowledging that while Enformer and Borzoi were trained on gene expression data, the gene expression tasks in `GUAN` in`E` are based on high-throughput dual reporter assays and have an esoteric distribution^3^ in a distinct organism, yeast. Differences in transcriptional mechanisms may affect performance in addition to the technical distribution shift induced by the dual reporter assay [10]. Nevertheless, the gpra-c and -d tasks from `GUAN` in`E` are included in our evaluation for completeness, and they offer an orthogonal view; one whose relative rankings surprisingly correlate with the field’s subjective notion of model ‘quality.’

#### Interpretable variable decomposition

Many sequence features are known to predict sequence function, GC-content chief among them. However, few interpretability techniques allow measuring a model’s *sensitivity* to easy-to-interpret features like GC-content, conservation, or chromosome size. We propose interpretable variable decomposition (IVD) as a solution to this issue.

While superficially similar to Shapley regression values [29], IVD is more closely related to first-order Sobol’ indices [50], and it has far less complexity than popular local explainability techniques [29, 47, 48, 31]. IVD is based on the approach proposed in Dinga et al. [12] and offers a superior runtime and holistic analysis (i.e. interfeature comparative analysis), unlike explainability aggregation techniques or computationally intense reverse-engineering approaches [47, 29]. However, in contrast to such introspective or perturbative techniques, it does require a significant amount of labeled data. This data is fairly rudimentary: a black box model’s predictions, a set of ground truth targets, and a set of features (endogenous or exogenous) for comparison, but the approach is less useful in scenarios without access to a moderate-scale benchmark or when suspected nonlinearities are major contributors to model sensitivity [50]. Given our use of the large-scale `GUAN` in`E` benchmark, IVD was a naturally advantageous choice that also allowed us to explore the underlying biology.

See the Methods section for more information about IVD and pseudocode, as well as an overview of our selected IVs. For readers interested in performing similar functional decompositions of variance, we would encourage seeking out additional novel featurizations and representations of biological mechanisms that can be used to evaluate genomics models.

## Results & Discussion

### Performance on the GUANinE benchmark

We report the few-shot performance scores of Enformer and Borzoi on `GUAN` in`E`’s test sets in Table 1 (along with previously reported baselines from [43]). Both models’ results are computed on the human output layer after an inverse softplus transform using an L2-regularized regression, as is common throughout large-scale AI evaluation [59, 7, 21]. Enformer’s results are the best of either single-bin or three-bin averages [1], while for a succint, yet comparable evaluation, we report Borzoi’s best performance across one, three, or five bin averages. Extended resolution analyses of Borzoi, up to 13 bin averaging (416 bp), are presented in Appendix Figure 13.

**Table 1.**
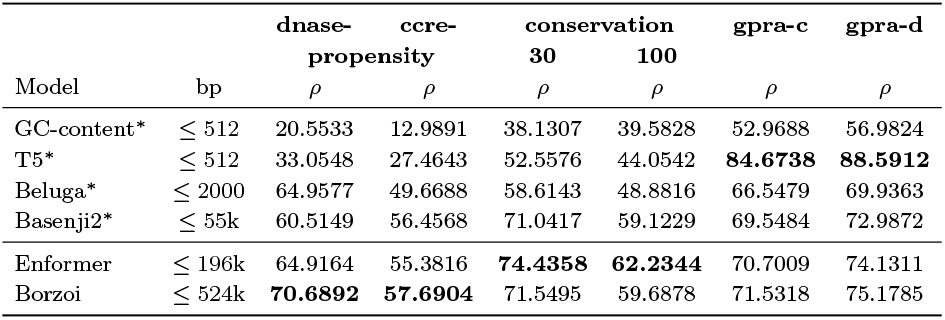
Performance of Enformer and Borzoi via linear evaluation on GUANinE. The T5 is the only fully-supervised model; other neural models are also few-shot (L2-regularized). Best score per task along with close scores (if any) are **bolded**. ^∗^Results reported previously [43].

### Embedding visualizations

#### Why intermediate embeddings?

While a model is trained and released with an output head attached, it is not uncommon for the ‘optimal’ representations to be found in the more information-dense intermediate layers, especially in biological language models [41]. For simplicity, we report performance using model outputs in Table 1, but we visualize and analyze single-bin final intermediate layer embeddings (3,072 and 1,920 dimensions for Enformer and Borzoi, respectively) for more compact representations^4^ with improved biological significance. We also report Enformer embedding performance in Appendix Table 2. We would advise readers aiming to maximize model performance to evaluate both options (outputs and embeddings).

#### Subjective Model Space

To better characterize the relationship between Enformer and Borzoi with `GUAN` in`E` tasks, we present a series of superimposed plots of `GUAN` in`E` sequences in the PCA space of Enformer’s training data embeddings in Figures 1 and 2. The former illustrates the distribution of the dnase-propensity task labels, colored by propensity (a correlate of regional DNase sensitivity). We observe that the unusual first PC of Enformer’s training dataset corresponds to a small number of inaccessible (and less conserved, see Appendix Figure 7) regions, as suggested by the predominantly class ‘0’ sequences from the dnase-propensity task. Concealed yet large-scale sequence variation is a likely explanation, although we were characterize these sequences. Endogenous retroviruses [32] were once such suspect we were unable to verify.

**Fig 1:**
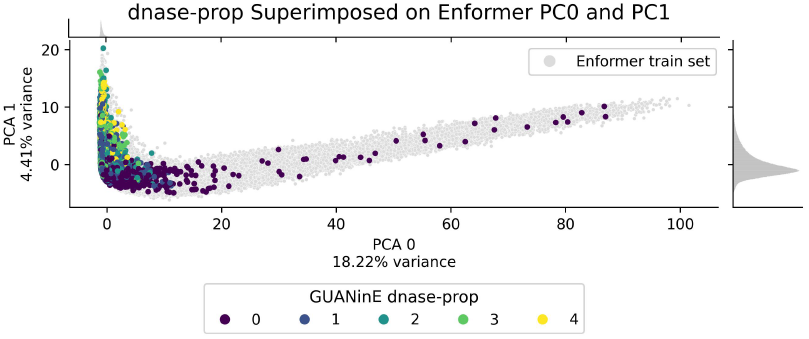
Enformer embedding PCA visualization of 1% of `GUAN` in`E`’s dnase-propensity examples (colored by propensity) superimposed on a grey background of 5% of Enformer’s training set sequences. The marginal density plots show that only a small number of sequences constitute the long tail of the ‘cirrus’ shape. A similar plot showing cons30 labels is presented in Appendix Figure 7.

**Fig 2:**
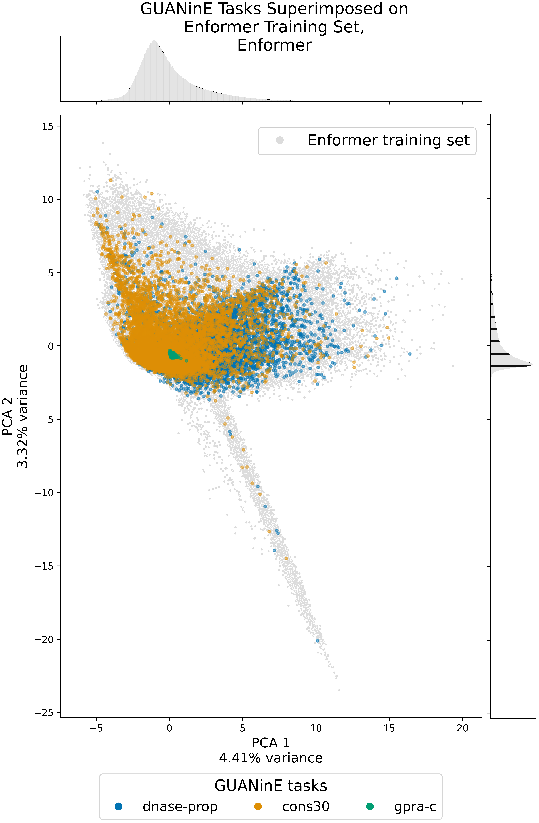
Enformer intermediate embedding PCA visualization of 1% of examples from select tasks of `GUAN` in`E` superimposed on a grey background of 5% of Enformer’s training set. `GUAN` in`E`’s dnase-propensity and cons30 tasks fall squarely within both models’ training distribution, while the gpra-c task is overly clustered near the origin. This suggests at minimum that these PCs, if not the models themselves, have failed to capture the variance of gpra-c task examples. A corresponding diagram for Borzoi on Enformer’s training set is located in Appendix Figure 6

Figure 2 illustrates how `GUAN` in`E` tasks mirror Enformer’s training distribution (comparable to Borzoi’s), except for the nearly singular distribution of gpra-c sequences. These cluster near the origin, suggesting the principal components (PCs) may not capture much variation in their embeddings. We investigated whether this visual lack of variation is due to a *statistical* lack of variance by computing the pairwise distances between embeddings, as visualized in Figure 3. The pairwise distance distributions, computed in the original, 3,072-dimensional embedding space of Enformer, confirm that the visual oversimilarity of gpra-c embeddings in PC-space is due to a lack of statistical variance; i.e the GPRA tasks are out-of-distribution, possibly because they are pushed towards an information-poor ‘null space’ of Enformer’s embedding space.

**Fig 3:**
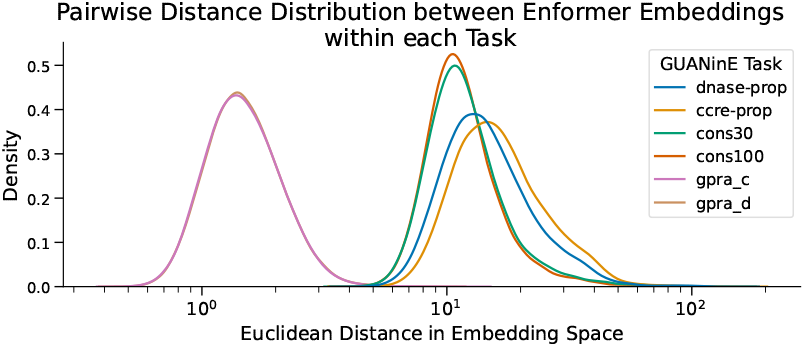
Pairwise distance KDE distributions, split by task, for single-bin embeddings from Enformer. For each task, we randomly sampled 100,000 pairs from 1% of the training set examples (<0.03% of all pairwise distances in the 1%). Both GPRA tasks (left, largely overlapping) have markedly lower average pairwise distances, suggesting little distinguishable difference between embeddings.

### Results of interpretable variable decomposition

The high-level findings of our variance explained decomposition are presented in Figure 4, where the proportion of variance explained unique to the deep learning models (blue) is separated from jointly explained variance (orange) using our enumerated sequence features (interpretable variables, IVs). The IVs we chose correspond to many basic sequence features, such as GC-content, as well as conservational and chromosomal features, as detailed further in Methods. In green (nearly invisible), we see the uncaptured variance explained by these exogenous, yet learnable, IVs. These negligible portions suggest that the models have absorbed the causal signal these factors have to offer, despite not receiving them during training.

**Fig 4:**
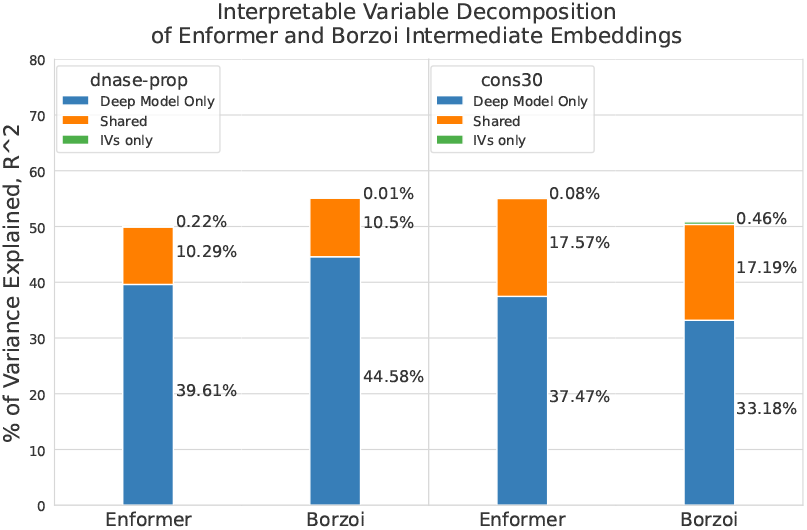
Percentage of variance explained by the deep learning models and by our enumerated sequence features (‘interpretable variables,’ IVs). The left two columns are for the dnase-propensity task, while the right two columns are the cons30 task. Not only can we quantify the utility of these IVs for each task, we can parse what information the deep models have captured beyond them. Notably, the ‘IVs only’ portion is small for most of the analyzed (model, task) pairs, implying that Enformer and Borzoi have subsumed these IVs entirely despite never seeing them during training.

For the dnase-propensity task, these interpretable variables explain up to 10.5% of the variance in linear contributions alone (before considering higher-order effects or interactions). Of that 10.5%, Enformer has surprisingly captured nearly 98% of it in the 10.3% jointly explained variance.

By analyzing the combined Enformer-IV model’s coefficients for the dnase-propensity task, we can measure the individual variables that contribute to this 10.5% of variance, as shown in Figure 5. Here, features are grouped by type (e.g. GC-content) and colored by scale (e.g. ‘core’ vs. ‘distal’, see Methods). The ‘core’ PhyloP30way conservation signal is by far the strongest predictor of dnase-propensity, but if it were not included, the ‘core’ GC-content would absorb most of its causal signal via omitted variable bias [14]. A similar visualization for the multiscale features used to decompose the cons30 task is presented in Appendix Figure 10.

**Fig 5:**
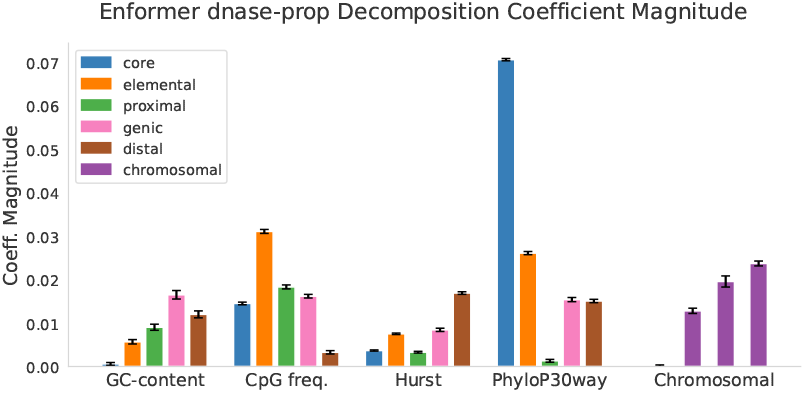
Coefficient magnitude (directionless) of the different features tested in the dnase-propensity IVD of Enformer’s predictions. All features were standardized to mean zero and variance one before coefficient estimation. Error bars shown here are 3.259 times the test-set estimated standard error for each coefficient; this number corresponds to the Bonferroni-corrected two-sided *p* < 0.01 threshold. `GUAN` in`E`’s large test set sample size is the reason coefficients can be estimated accurately despite partial collinearity. Matching tables for cons30 and Borzoi are located in the Appendix

### Models incorporate conservation in their predictions

We find that Enformer is a strong predictor of sequence conservation (cons30 and cons100) (Table 1), whose connection to deleteriousness and pathogenicity suggests that Enformer may be relevant for medical and clinical applications, at least where region-level conservation signal is important. Interestingly, Borzoi has slightly lower performance in predicting conservation, suggesting that while it has additional context to leverage, it may not fully utilize relevant sequence information at intermediate scales (within Enformer’s resolution). Enformer’s greater model depth is a likely explanation for this discrepancy, given the difficulty of estimating conservation from sequence alone [43].

We suspect that this understanding of conservation contributes to Enformer’s performance on other tasks – for example, compared to Basenji2 [1], which undergoes the same training procedure, Enformer shows considerably higher performance in identifying DNase-hypersensitive regions (dnase-propensity, Table 1). Notably, this improvement does not transfer into tissue-specific *functional* understanding, as seen by Basenji2’s superior ccre-propensity performance. Succeeding on `GUAN` in`E`’s ccre-propensity task requires accurately estimating sequence functionality (i.e. histone marks) conditioned on a region being hypersensitive to DNase.

As our interpretable variable decomposition shows, Enformer misses out on only a tiny portion (0.08%, third column, Figure 4) of variance explainable by the evaluated exogenous IVs. To achieve such low IVs-only scores, both Enformer and Borzoi have learned the implicit primary DNA sequence signatures of chromosomal features, which are known to impact sequence conservation [58].

### What do larger models add?

While Enformer outperforms Basenji2 on most of the tested tasks in `GUAN` in`E` , it lags behind in its ability to recognize cis-regulatory element propensity (i.e. histone marks). It also merely matches Beluga in dnase-propensity, despite Beluga’s substantially smaller context size of 2,000 bp. Conversely, Borzoi lags substantially behind Enformer in understanding sequence conservation. Despite this shortcoming, it is better at both identifying accessible regions (dnase-propensity) and understanding their function (ccre-propensity).

Enformer and Borzoi may be optimized for different objectives than accessibility or conservation, respectively, e.g. gene expression performance (CAGE) and RNA-seq, cumulatively. It is worth recognizing that the areas that any given model “outperforms” on may be offset by decreased performance on other tasks, especially in a multitask regime [51]. Clearly, there is a need for the field to recognize that no one model can succeed at all tasks – and perhaps building smaller, more specialized genomic models may be more productive in many cases.

### Visualization insights into generalizability

A key insight enabled by our framework is Enformer’s and Borzoi’s limited generalizability to the out-of-distribution GPRA tasks in `GUAN` in`E` , even with few-shot learning. By visualizing their embeddings, we observe their tendency to map the random oligonucleotide sequences to an information-poor ‘null space,’ and it stands to reason that perhaps straightforward embedding-space-based solutions like adapter layers [18] or domain adaptation [4] may be able to improve their performance.

An interesting potential genomics-specific form of transfer learning may be through the use of homology information, perhaps in a manner leveraging cross-organismal relationships in well-studied model organisms [6]. This approach could be evaluated by fine-tuning models like Enformer on such a large dataset as the GPRA tasks.

### Interpretability untangles biological function

The IVD analysis we undertake is complementary to motif discovery [48] or gene annotation, and we would encourage readers interested in interpretability to explore such featurizations.

Using our IVD approach, we can conclude that not only has Enformer learned to leverage sequence conservation and chromosomal features (e.g. telomeric distance), but also that these features are intrinsically embedded *in primary sequence alone*, as Enformer does not receive additional inputs. This finding has profound implications in terms of how we choose to discuss DNA accessibility (i.e. relative to chromosomal position), and in terms of what factors belie a model’s test set performance and the physical assays we use to collect data [10, 54]. We leave to future work how genomics researchers can best leverage or de-confound these signals.

#### Learning more than deep learning

If genomics researchers are able to articulate the explanatory variables underlying a deep learning model’s success (be it GC-content, conservation, or chromosomal features, etc.), this would reduce the need to rely upon inefficient^5^ feature extraction approaches dependent on deep neural networks. Additionally, enumerating these interpretable features can assist us to debug and improve existing neural models by indicating whether a model has already learned a potential ‘ensemble’ feature [22, 19]. Our analysis here indicates that augmenting Enformer with *region*-level conservation scores and simplistic chromosomal features would not improve performance on downstream tasks, as the model has already subsumed their signal, but that Borzoi could marginally benefit from such features, as evidenced by the 0.45% IV-only portion of variance explained in the rightmost column of Figure 4.

#### Conservation as an indicator of sequence function

One puzzling finding from our interpretation analysis relates to Enformer being a strong predictor of sequence conservation (Table 1), although ≈ 45% of variance in PhyloP30way conservation is uncaptured by Enformer (third column, Figure 4). Yet surprisingly, this untapped portion of conservation adds at most a marginal (0.22% of variance, leftmost column, Figure 4) amount of signal to the dnase-propensity task. Of course, we are merely examining *linear* relationships of conservation to dnase-propensity, and more exhaustive measures of sensitivity may reveal nonlinear or higher-order effects [50].

We hypothesize that Enformer in particular arrives at a “conservation-aware embedding space” which captures the portion of sequence conservation relevant to its downstream tasks. This is likely related to the utility of sequence conservation for discovering functional elements [38, 49], as many yet unexplored sequence features (coding regions, binding motifs, and splice junctions, etc.) may serve as proxies for both accessibility and conservation for Enformer and Borzoi during training. Borzoi, on the other hand, may not outperform Enformer in all aspects without greater model depth (e.g. cons30 and cons100), though it remains the best model evaluated on `GUAN` in`E`’s accessibility (dnase-propensity) and accessible region functionality (ccre-propensity) tasks.

### Reflections on GUANinE

Finally, we discuss a handful of insights regarding benchmarking that have emerged through this case study. Current models such as Enformer exhibit high performance on some tasks, but are far from having solved biology *in general* – or from necessarily being the proper modeling approach to do so. Given that `GUAN` in`E`’s propensity tasks benefit from aggregation across cell lines [43] (thus reducing the proportion of experimental noise, due to the law of large numbers), we believe the performance ceiling on these tasks is far from being reached. Furthermore, models like Enformer, which have been trained on a superset of `GUAN` in`E`’s propensity tasks [43], should be able to achieve near perfect accuracy on dnase- and ccre-propensity, as these tasks are straightforward derivations of such data. The fact that models like Enformer explain just half of the variance in these tasks indicates that these models have not fully learned their own training data.

A key limitation of `GUAN` in`E` for evaluating a model like Enformer on gene expression performance is its reliance on GPRA data for expression (to circumvent the small-N and collinearity of human expression data). Models like Enformer struggle to perform on personal gene expression estimation [20], but we do not believe the gpra-c or gpra-d tasks in `GUAN` in`E` currently illustrate that issue. As is, they instead show the limitation of models like Enformer when attempting to predict gene expression on exogenous sequences and different organisms without additional fine-tuning – and they offer a rich opportunity for innovation in transfer learning techniques.

An important follow-up to our analysis entails whether *regional* conservation and accessibility is transferable to finer-grained resolutions [2] or variant-interpretation tasks, for which some evidence of non-performance exists [20]. While many researchers are interested in applying sequence models to precision genomics tasks, it is unknown how useful accessibility detection or regulatory element recognition are for precision medicine, despite these tasks being the vest majority of training data for supervised CNN models like Enformer or Borzoi [61, 62, 26, 2, 25, 1].

## Methods

Notably, we do not perform any fine-tuning of Enformer or Borzoi, relying instead on an L2-regularized linear evaluation fit to model outputs. Coefficients, formulae, and details regarding the runtime and feature extraction are in the Appendices.

### Interpretable Variable Decomposition

For both the dnase-propensity and cons30 tasks we decompose the variance explained into enumerated (“interpretable variable”) and unenumerated (unspecified) portions of our few-shot regression models on Enformer outputs. This is accomplished via the post-hoc deconfounding analysis methodology proposed in Dinga et al. [12], which relies on the Pythagorean properties of the RSS (residual sum of squares), in canonical-link linear models [17]. This functional decomposition of variance is loosely related to the explainability application of game-theoretic Shapley values [29, 47], but it instead focuses on global interpretation, much like first-order Sobol’ indices [50].

#### IVD Algorithm

Our procedure is as follows:

1. Fit combined linear model *f*_*IV* ∪*E*_ (*X*_train_) = *y*_train_ on both interpretable variables *IV* and model predictions *E* (the latter being the L2-regularized few-shot prediction validated on `GUAN` in`E`’s development set, although any predictor will do).
2. Fit reduced models *f*_*IV*_ and *f*_*E*_ as before.
3. Determine the RSS and R^2^ of *f*_*IV* ∪*E*_, *f*_*IV*_ , and *f*_*E*_ on an independent (and thus unbiased) test set {*X*_test_, *y*_test_}
4. Adjust for the partial correlation effects of the *R*^2^ measures as in Dinga et al. [12].
5. Derive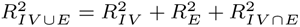

Notably, it does not require the use of heuristics or approximations, access to model parameters, nor expensive computational resources (all typically necessary for global methods based on aggregating local explainability [31]).

#### Limitations to IVD

In practice, three issues can arise when attributing variance to specific coefficients or to reduced models as we do here – we briefly address all three below.

First, as discussed in Hastie [17], there is a bias on coefficient estimates due to overfitting when performing ordinary least squares (OLS). This is easily resolved by instead estimating RSS on an independent test set, i.e. step 3 in our IVD algorithm above, as discussed in Dinga et al. [12].

Second, as IVs are almost certainly correlated with model predictions (many are even literal *confounders*), there is a micronumerosity problem, i.e. sufficient sample size is required to reliably estimate individual coefficients in the presence of partial collinearity [16]. This is of limited concern due to our large (*N >* 40000) sample sizes in `GUAN` in`E` , and we report all p-values, coefficients, and standard errors in Appendix C. This issue is not dissimilar to the partial collinearity resulting from the application of linear models to GWAS or fine-mapping.

Third, the predominant – and underappreciated – limitation to IVD is *omitted variable bias* [14]. Specifically, if an identifiable, yet *un*included, covariate is correlated with a *proxy* variable in our regression model, the explanatory power of this absentee variable will be absorbed by its correlates (proxies). This will inevitably inflate our estimates of variance explained by the specific variables under investigation. To counteract this, we select multiple redundant features for each regression, and within the dnase-propensity regression, we specifically include mean PhyloP30way scores to prevent GC-content and other variables from absorbing their signal.

PhyloP30way scores are not included as IVs for the cons30 task to avoid circularity.

### Multiscale features for the decomposition

Numerous biologically endogenous and exogenous sequence features have been associated with DNA function in multivariate analysis [36, 37, 23]. The proportion of guanine plus cytosine content (G+C or GC-content) is among the simplest yet most predictive in many organisms [53]. For our variance explained decomposition regressions described in subsection 4.1, we enumerate a number of both sequence-level and chromosome-level features measured at multiple scales.

#### Sequence-level features

We include GC-content, CpG frequency, and the Hurst coefficient of alternating Purine-Pyrimidine content [36, 37] at each choice of scale. We also regressed DNA melting points but observed that melting point estimates correlate perfectly with GC-content, as in Biopython’s libraries for DNA melting points [8]. The scales of analysis chosen for these features were 127bp, 511bp, 2047bp, 8191bp, and 32767bp, referred to as ‘core’, ‘element’, ‘proximal’, ‘genic’, and ‘distal’, respectively.

For each (scale, feature) combination, the measurement was centered to the example, then the corresponding measurement from the next-smallest region was subtracted. For example, the ‘genic_gc’ feature measures GC-content of only the ≈ 6kbp surrounding the ‘proximal’ sequence. This nested featurization provides partially decorrelated features, reducing variance inflation. For the dnase-propensity regression, the mean PhyloP30way inputs are calculated similarly.

#### Chromosome-level features

Unlike sequence features which operate directly on a local scale, numerous chromosome-level background features influence a chromosome’s constituent sequences, e.g. smaller chromosomes having increased linkage and background conservation due to failed recombination events [58]. We choose to include a feature set that is both overdetermined yet rich: the log of the hg38 chromosome size, the distance to the arm’s telomere (in bp), the relative position of a sequence in the chromosome’s arm (i.e. 0.0 to 1.0, where 0.5 is midway between centromere and telomere), and the distance to the nearest centromere (in bp). The chromosomal feature coefficients included in Figure 5 are in the same order.

All missing values (e.g. unalignable regions for phyloP) were imputed to the mean, and we standardize all features to mean of zero and variance of one to aid with model fitting and coefficient interpretation.

## Conclusion

Through our Enformation Theory framework of performance, visualization, and interpretation, we arrive at a deeper understanding of a model’s performance than metrics reporting alone. We discuss the advances and limitations of recent large-context models on the `GUAN` in`E` benchmark and identify *what* types of interpretable variables and sequence features they have learned to achieve such performance. We show, with significance, their reliance on information contained in features such as conservation and distance to centro- and telomeric regions, and we highlight the need to better understand the role of these features in experimental measurements and model predictions. We illustrate a key generalization issue of models like Enformer and Borzoi to synthetic GPRA sequences and affirm the role of `GUAN` in`E` as a leading benchmark for genomic AI models. It is our hope that the analyses presented in this framework, such as our use of interpretable variable decomposition to identify biologically meaningful covariates, will become more ubiquitous throughout the field of genomic AI and supply it with a unified toolkit for model evaluation.

## Acknowledgements

We thank Ruchir Rastogi and other members of the Ioannidis lab for invaluable feedback. Special thanks to Amelia Dobis for her assistance in copyediting. This work was funded in part by grants from the UC Noyce Initiative for Computational Transformation and the Chan Zuckerberg Biohub San Francisco.

## Code Availability

The roughly ∼ 60 GB of Borzoi model outputs and ∼ 40 GB of Enformer model outputs used to generate the best dev-set-performance results seen in Table 1 will be made available at https://github.com/ni-lab/enformation-theory. We will also include intermediate model embeddings and code for replicating our IVD analysis, along with instructions for recreating PCA figures.

## Runtime and Extended Methods

### Feature extraction with Enformer

We use the PyTorch [33], variable-length implementation of Enformer^6^ for inference efficiency. For each task in `GUAN` in`E` , we align inputs to the center-left bin of Enformer (the 448th bin, or 447 when zero-based indexing), and we provide the full 196kbp + 256bp of sequence context. This allows us to access both the single bin embeddings and prediction for each target as well as a three-bin average. Incomplete sequences extending into telomeric regions were padded with `N` Ns, although we do not see a strong association between `N` Ns and Enformer embeddings, see Appendix 9.

The 3,072-dimensional embeddings from Enformer, three-bin-averaged and single-bin, along with the 5313-dimensional outputs from the human prediction head, three-bin-averaged and single-bin, were cached for our linear evaluation step. These embeddings and outputs were computed for the full development and test sets of each task in `GUAN` in`E` , along with a 1% subsample of the training sets provided in `GUAN` in`E`’s Hugging Face repository^7^.

All features from the output layer were passed into an inverse-softplus transformation, which slightly improved performance in every task (likely due to the softplus destroying ratio information).

### Linear evaluation

We use scikit-learn’s [35] ridge regression to perform L2-regularized linear evaluation on the three-bin-averaged and single-bin embeddings and outputs from Enformer. We searched over regularization coefficients of [0.1, 0.3, 1., 3., 3.e5, 1.e6], evaluated on the development split, as we noticed substantial variability in optimal regularization coefficients across tasks (many of the regressions required high amounts of regularization, possibly due to the combination of high dimensionality and collinearity). The best of either the three-bin-averaged or single-bin predictions on the development set was used for test set predictions, as reported in Table 1.

### Dimensionality reduction

Due to the large size of Enformer’s training set, we use `incremental PCA` [45, 35] to estimate the first 30 principal components without resorting to subsetting. For all visuals, dimension reduction was performed on the single-bin embeddings. The 30 PCs explained a combined 51.6122% of Enformer’s training set variance.

`t-SNE` (in the Appendices) was performed on examples after first being reduced to 30 PCs for runtime efficiency.

## Additional figures

**Table 2.**
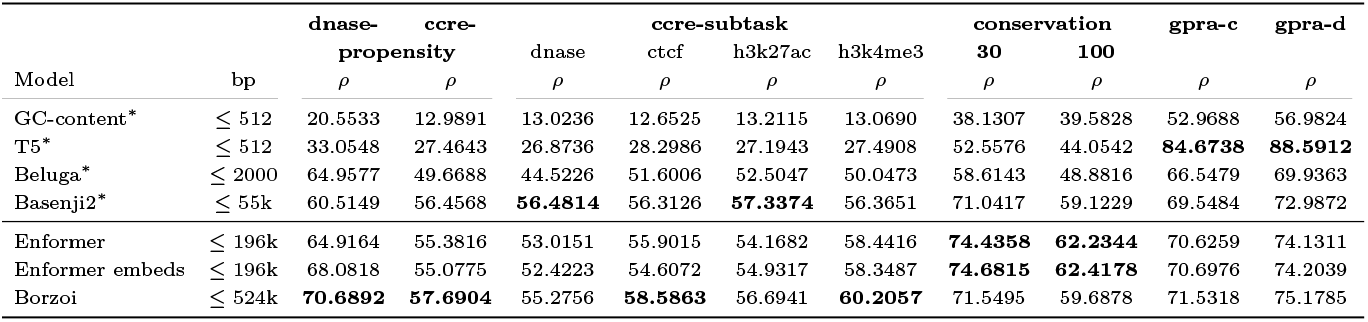
Performance of Enformer’s outputs and embeddings via linear evaluation on GUANinE. The T5 is a fully-supervised model while other neural models are also few-shot (L2-regularized), as Enformer. Best score per task along with close scores (if any) are **bolded**. ^∗^Non-Enformer results reported previously [43]. Single-bin embeddings results.

**Fig 6:**
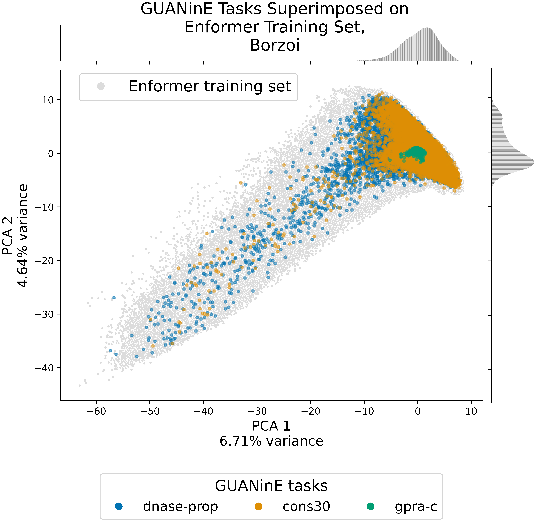
Borzoi intermediate embedding PCA visualization of 1% of examples from select tasks of `GUAN` in`E` superimposed on a grey background of 5% of Enformer’s training set. `GUAN` in`E`’s tasks are better distributed in Borzoi embeddings compared to Enformer’s, and the gpra-c task, while clustered, has some shape, which matches its improved performance compared to Enformer.

**Fig 7:**
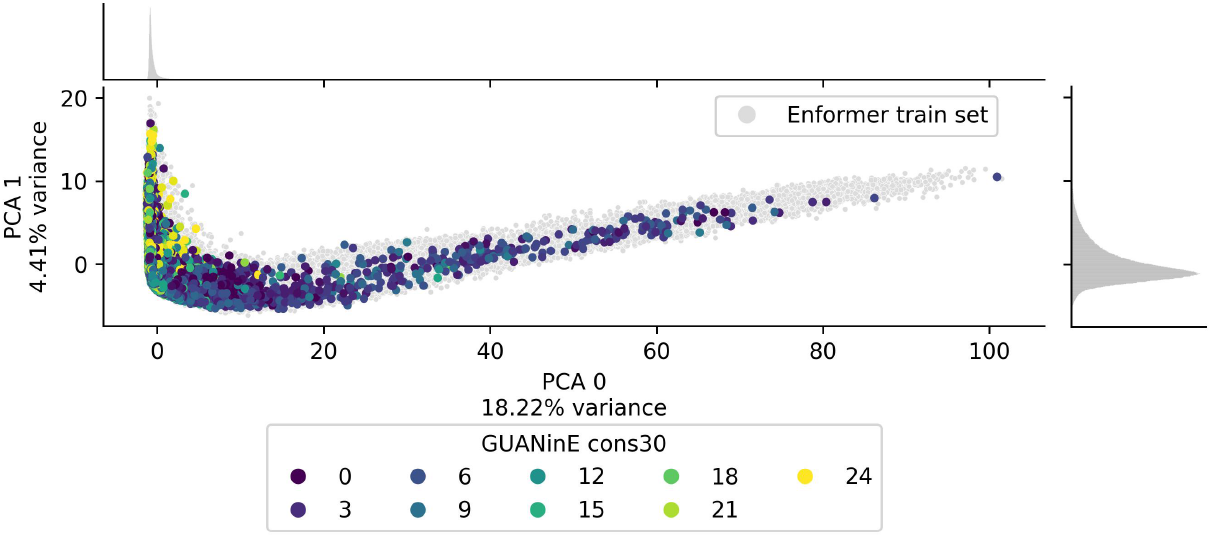
Enformer embedding PCA visualization of 1% of `GUAN` in`E`’s cons30 examples (colored by binned conservation label) superimposed on a grey background of 5% of Enformer’s training set sequences. The marginal density plots show that only a small number of sequences constitute the long tail of the ‘cirrus’ shape, which is enriched for unconserved (closer to zero) values.

**Fig 8:**
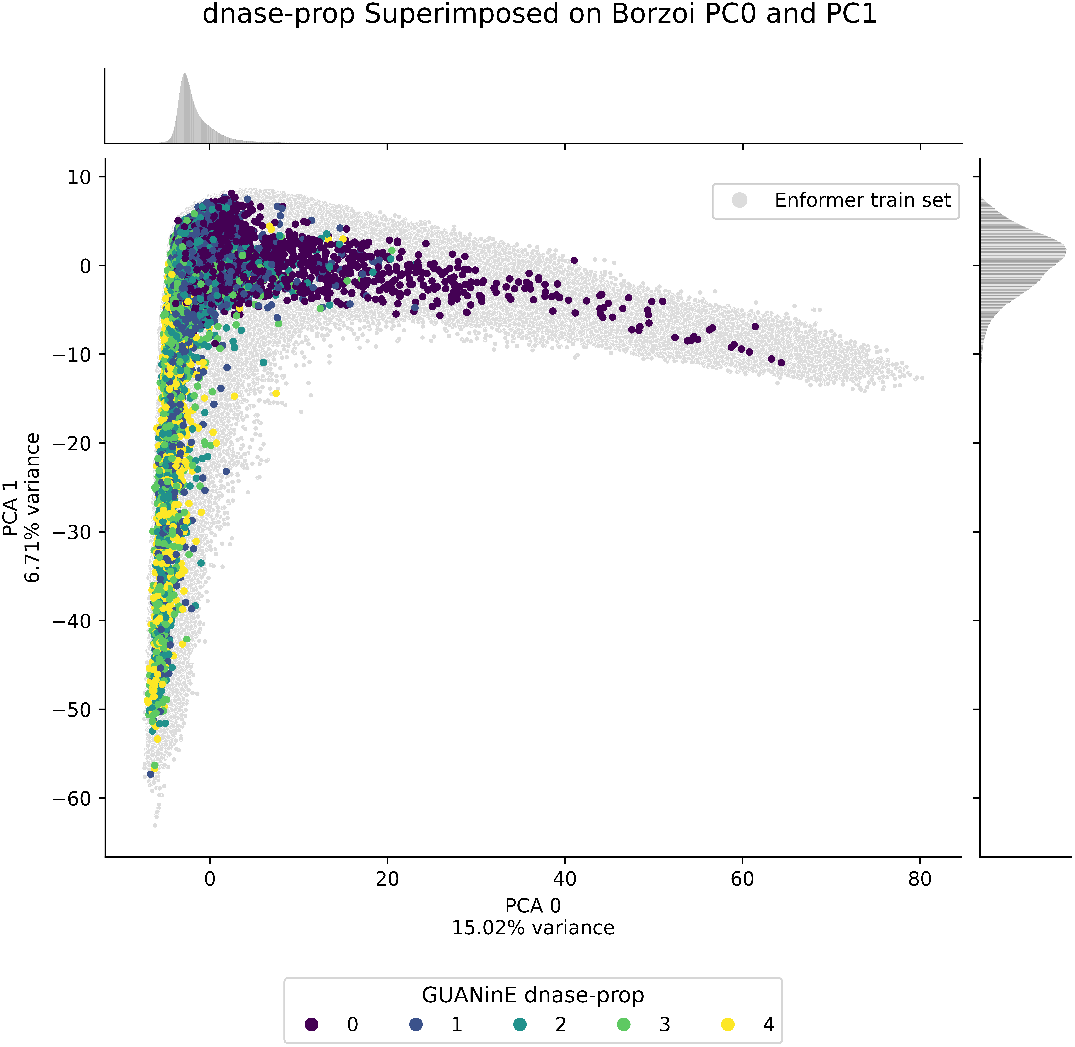
Borzoi embedding PCA visualization of 1% of `GUAN` in`E`’s cons30 examples (colored by binned conservation label) superimposed on a grey background of 5% of Enformer’s training set sequences. The ‘boomerang’ shape is not as pronounced as Enformer’s, due partly to the difference in explained variance per component, although the distribution is highly similar.

**Fig 9:**
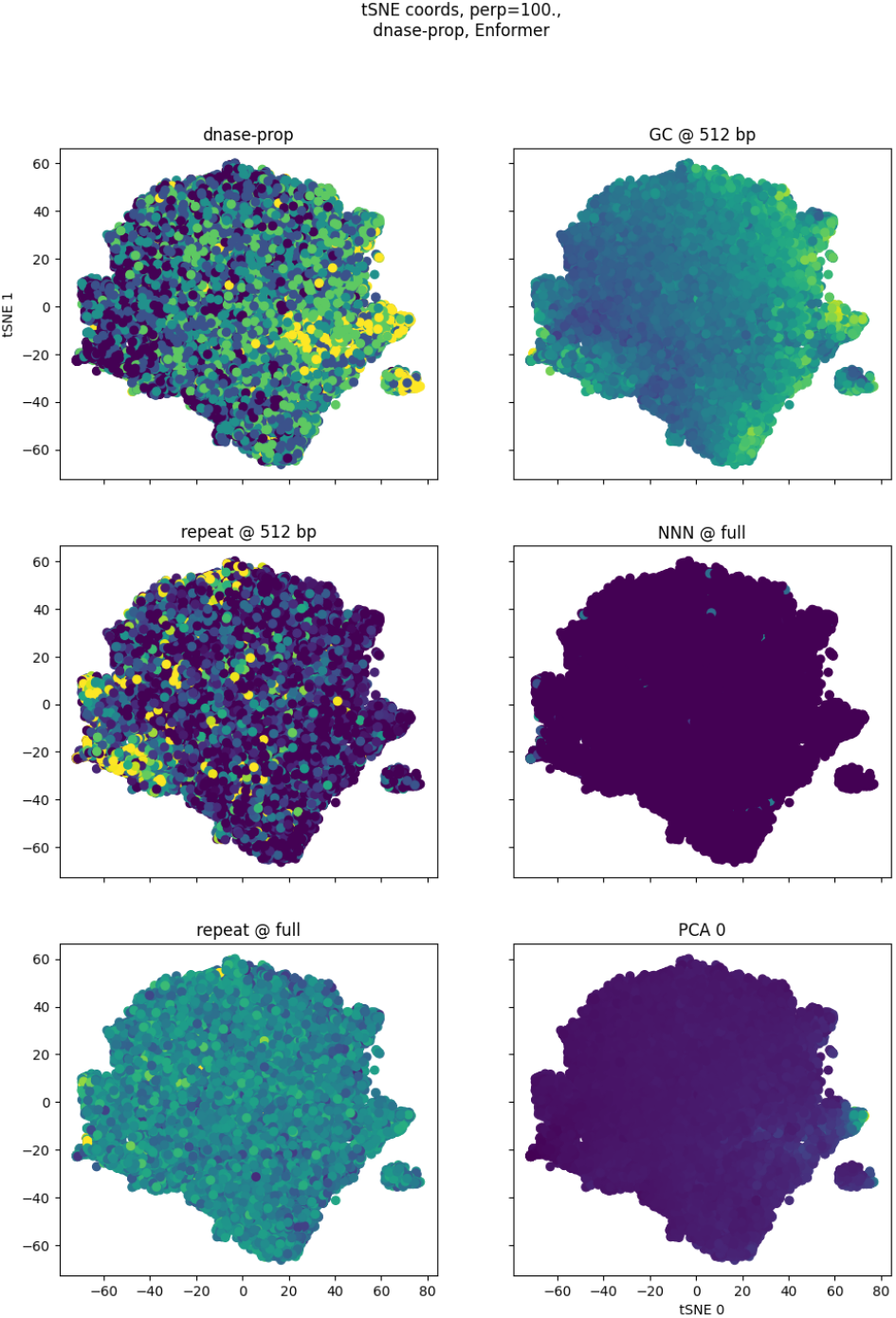
`t-SNE` of Enformer embeddings of examples from the dnase-propensity task. Clockwise from the top left, we visualize dnase-propensity scores (yellow is ‘4’, purple is ‘0’); GC-content of the ‘element’ region at 512 bp; repeat frequencies of the ‘element’ region, as determined by RepeatMaster (yellow is greater); the number of Ns at the full context size (yellow is greater); the repeat proportion at the full context size; and the first PCA component.

## Enformer coefficients and test statistics for the combined model regressions

**Table 3.**
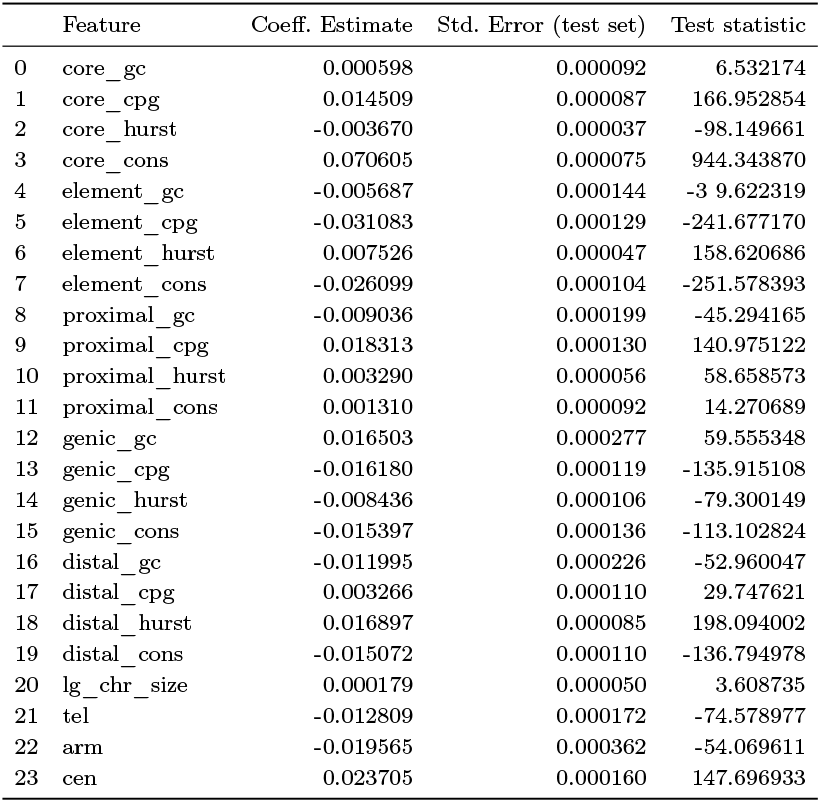
Combined model regression for the IVD on the dnase-propensity task estimates of Enformer. This regression is fit to Enformer’s intermediate representation embeddings, as discussed in the Methods section. The degrees of freedom for involving the standard error, which we complete on the test set, is 44801 − 24 − 3072 = 41705, which renders the use of z-scores appropriate.

|  | Feature | Coeff. Estimate | Std. Error (test set) | Test statistic |
| --- | --- | --- | --- | --- |
| 0 | core_gc | 0.000598 | 0.000092 | 6.532174 |
| 1 | core_cpg | 0.014509 | 0.000087 | 166.952854 |
| 2 | core_hurst | -0.003670 | 0.000037 | -98.149661 |
| 3 | core_cons | 0.070605 | 0.000075 | 944.343870 |
| 4 | element_gc | -0.005687 | 0.000144 | -3 9.622319 |
| 5 | element_cpg | -0.031083 | 0.000129 | -241.677170 |
| 6 | element_hurst | 0.007526 | 0.000047 | 158.620686 |
| 7 | element_cons | -0.026099 | 0.000104 | -251.578393 |
| 8 | proximal_gc | -0.009036 | 0.000199 | -45.294165 |
| 9 | proximal_cpg | 0.018313 | 0.000130 | 140.975122 |
| 10 | proximal_hurst | 0.003290 | 0.000056 | 58.658573 |
| 11 | proximal_cons | 0.001310 | 0.000092 | 14.270689 |
| 12 | genic_gc | 0.016503 | 0.000277 | 59.555348 |
| 13 | genic_cpg | -0.016180 | 0.000119 | -135.915108 |
| 14 | genic_hurst | -0.008436 | 0.000106 | -79.300149 |
| 15 | genic_cons | -0.015397 | 0.000136 | -113.102824 |
| 16 | distal_gc | -0.011995 | 0.000226 | -52.960047 |
| 17 | distal_cpg | 0.003266 | 0.000110 | 29.747621 |
| 18 | distal_hurst | 0.016897 | 0.000085 | 198.094002 |
| 19 | distal_cons | -0.015072 | 0.000110 | -136.794978 |
| 20 | lg_chr_size | 0.000179 | 0.000050 | 3.608735 |
| 21 | tel | -0.012809 | 0.000172 | -74.578977 |
| 22 | arm | -0.019565 | 0.000362 | -54.069611 |
| 23 | cen | 0.023705 | 0.000160 | 147.696933 |

**Table 4.**
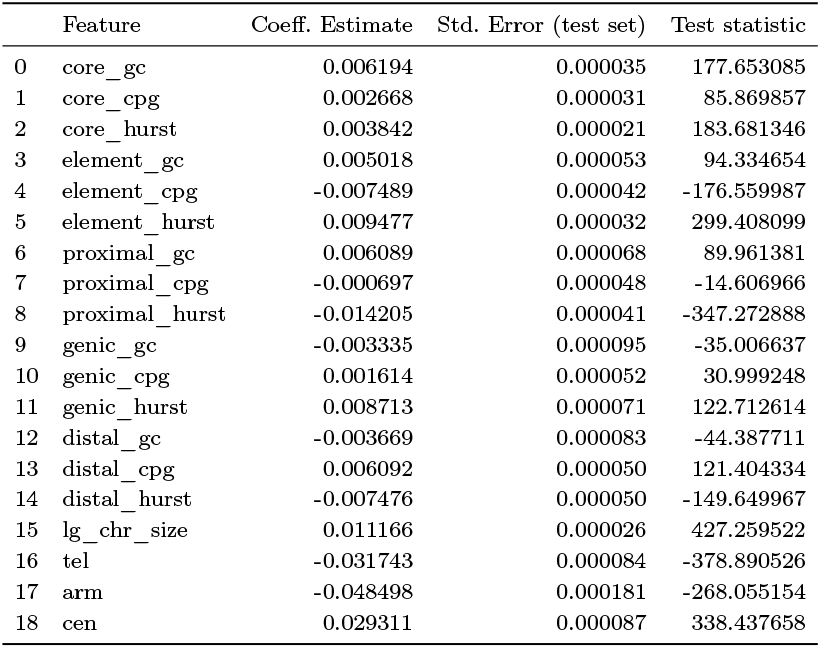
Combined model regression for the IVD on the cons30 task estimates of Enformer. This regression is fit to Enformer’s intermediate representation embeddings, as discussed in the Methods section. The degrees of freedom for involving the standard error, which we complete on the test set, is 44801 − 19 − 3072 = 41710, which renders the use of z-scores appropriate.

|  | Feature | Coeff. Estimate | Std. Error (test set) | Test statistic |
| --- | --- | --- | --- | --- |
| 0 | core_gc | 0.006194 | 0.000035 | 177.653085 |
| 1 | core_cpg | 0.002668 | 0.000031 | 85.869857 |
| 2 | core_hurst | 0.003842 | 0.000021 | 183.681346 |
| 3 | element_gc | 0.005018 | 0.000053 | 94.334654 |
| 4 | element_cpg | -0.007489 | 0.000042 | -176.559987 |
| 5 | element_hurst | 0.009477 | 0.000032 | 299.408099 |
| 6 | proximal_gc | 0.006089 | 0.000068 | 89.961381 |
| 7 | proximal_cpg | -0.000697 | 0.000048 | -14.606966 |
| 8 | proximal_hurst | -0.014205 | 0.000041 | -347.272888 |
| 9 | genic_gc | -0.003335 | 0.000095 | -35.006637 |
| 10 | genic_cpg | 0.001614 | 0.000052 | 30.999248 |
| 11 | genic_hurst | 0.008713 | 0.000071 | 122.712614 |
| 12 | distal_gc | -0.003669 | 0.000083 | -44.387711 |
| 13 | distal_cpg | 0.006092 | 0.000050 | 121.404334 |
| 14 | distal_hurst | -0.007476 | 0.000050 | -149.649967 |
| 15 | lg_chr_size | 0.011166 | 0.000026 | 427.259522 |
| 16 | tel | -0.031743 | 0.000084 | -378.890526 |
| 17 | arm | -0.048498 | 0.000181 | -268.055154 |
| 18 | cen | 0.029311 | 0.000087 | 338.437658 |

## Borzoi coefficients and test statistics for the combined model regressions

**Table 5.** Combined model regression for the IVD on the dnase-propensity task estimates of Borzoi. This regression is fit to Borzoi’s intermediate representation embeddings, as discussed in the Methods section. The degrees of freedom for involving the standard error, which we complete on the test set, is 44801 − 24 − 1920 = 41857, which renders the use of z-scores appropriate.

|  | Feature | Coeff. Estimate | Std. Error (test set) | Test statistic |
| --- | --- | --- | --- | --- |
| 0 | core_gc | 0.005392 | 0.000092 | 58.351982 |
| 1 | core_cpg | 0.004118 | 0.000088 | 46.754103 |
| 2 | core_hurst | -0.001680 | 0.000038 | -44.333784 |
| 3 | core_cons | 0.058776 | 0.000076 | 773.763469 |
| 4 | element_gc | -0.001803 | 0.000146 | -12.388969 |
| 5 | element_cpg | -0.011611 | 0.000130 | -89.083741 |
| 6 | element_hurst | 0.005318 | 0.000048 | 110.581207 |
| 7 | element_cons | -0.014394 | 0.000105 | -136.957991 |
| 8 | proximal_gc | 0.010133 | 0.000202 | 50.079303 |
| 9 | proximal_cpg | 0.000205 | 0.000132 | 1.553287 |
| 10 | proximal_hurst | 0.005928 | 0.000057 | 104.271313 |
| 11 | proximal_cons | -0.012964 | 0.000093 | -139.343436 |
| 12 | genic_gc | 0.001663 | 0.000281 | 5.920405 |
| 13 | genic_cpg | -0.001510 | 0.000121 | -12.518792 |
| 14 | genic_hurst | -0.011763 | 0.000108 | -109.115536 |
| 15 | genic_cons | -0.000539 | 0.000138 | -3.902648 |
| 16 | distal_gc | -0.020875 | 0.000230 | -90.928839 |
| 17 | distal_cpg | 0.011651 | 0.000111 | 104.678594 |
| 18 | distal_hurst | 0.014653 | 0.000086 | 169.491340 |
| 19 | distal_cons | -0.017507 | 0.000112 | -156.776356 |
| 20 | lg_chr_size | 0.008811 | 0.000050 | 175.143389 |
| 21 | tel | -0.017736 | 0.000174 | -101.896018 |
| 22 | arm | -0.029773 | 0.000367 | -81.200866 |
| 23 | cen | 0.026073 | 0.000163 | 160.321390 |

**Table 6.**
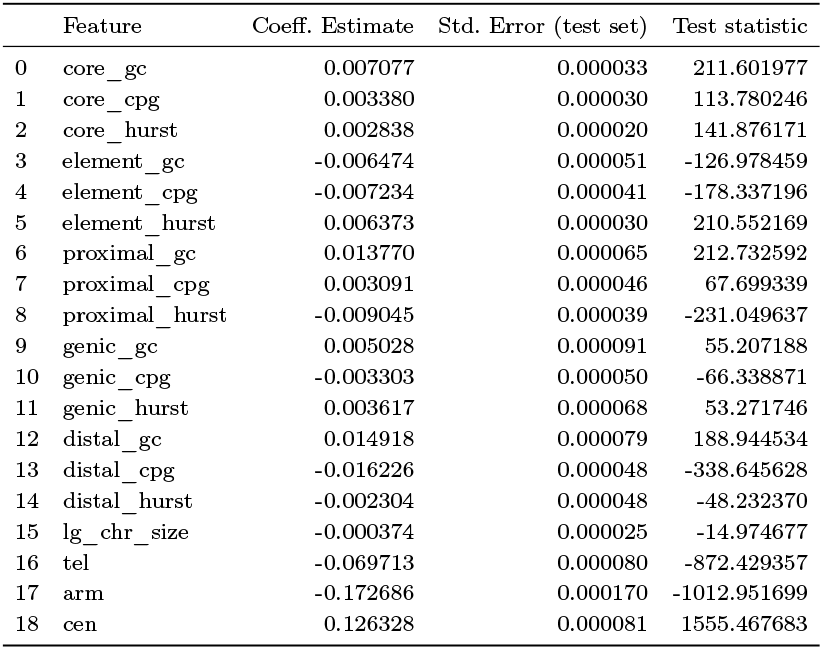
Combined model regression for the IVD on the cons30 task estimates of Borzoi. This regression is fit to Borzoi’s intermediate representation embeddings, as discussed in the Methods section. The degrees of freedom for involving the standard error, which we complete on the test set, is 44801 − 19 − 1920 = 41862, which renders the use of z-scores appropriate.

|  | Feature | Coeff. Estimate | Std. Error (test set) | Test statistic |
| --- | --- | --- | --- | --- |
| 0 | core_gc | 0.007077 | 0.000033 | 211.601977 |
| 1 | core_cpg | 0.003380 | 0.000030 | 113.780246 |
| 2 | core_hurst | 0.002838 | 0.000020 | 141.876171 |
| 3 | element_gc | -0.006474 | 0.000051 | -126.978459 |
| 4 | element_cpg | -0.007234 | 0.000041 | -178.337196 |
| 5 | element_hurst | 0.006373 | 0.000030 | 210.552169 |
| 6 | proximal_gc | 0.013770 | 0.000065 | 212.732592 |
| 7 | proximal_cpg | 0.003091 | 0.000046 | 67.699339 |
| 8 | proximal_hurst | -0.009045 | 0.000039 | -231.049637 |
| 9 | genic_gc | 0.005028 | 0.000091 | 55.207188 |
| 10 | genic_cpg | -0.003303 | 0.000050 | -66.338871 |
| 11 | genic_hurst | 0.003617 | 0.000068 | 53.271746 |
| 12 | distal_gc | 0.014918 | 0.000079 | 188.944534 |
| 13 | distal_cpg | -0.016226 | 0.000048 | -338.645628 |
| 14 | distal_hurst | -0.002304 | 0.000048 | -48.232370 |
| 15 | lg_chr_size | -0.000374 | 0.000025 | -14.974677 |
| 16 | tel | -0.069713 | 0.000080 | -872.429357 |
| 17 | arm | -0.172686 | 0.000170 | -1012.951699 |
| 18 | cen | 0.126328 | 0.000081 | 1555.467683 |

## Additional coefficient magnitude figures

**Fig 10:**
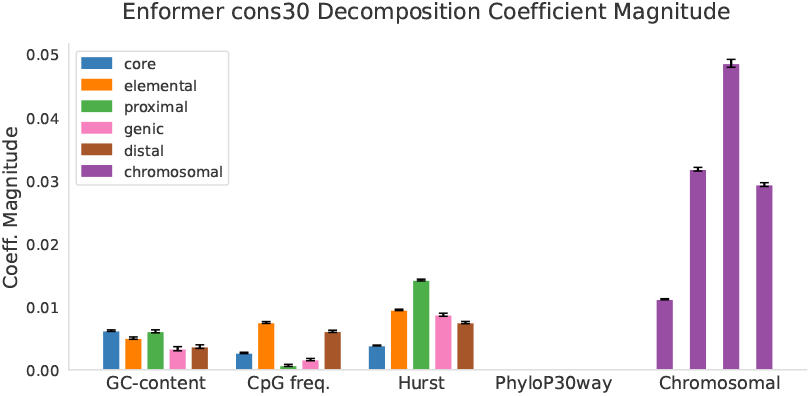
Coefficient magnitude (directionless) of the different features tested in the cons30 IVD of Enformer’s predictions. All features were standardized to mean zero and variance one before coefficient estimation. Error bars shown here are 3.259 times the test-set estimated standard error for each coefficient; this number corresponds to the Bonferroni-corrected two-sided *p* < 0.01 threshold. To prevent circularity issues, PhyloP30way features were not used here (features align vertically).

**Fig 11:**
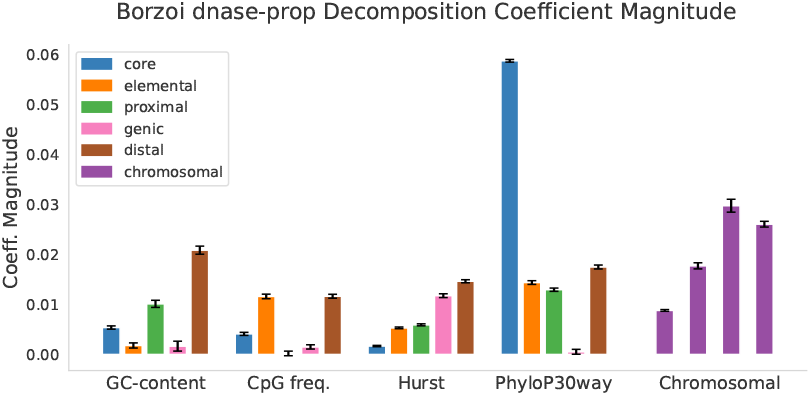
Coefficient magnitude (directionless) of the different features tested in the dnase-propensity IVD of Borzoi’s predictions. All features were standardized to mean zero and variance one before coefficient estimation. Error bars shown here are 3.259 times the test-set estimated standard error for each coefficient; this number corresponds to the Bonferroni-corrected two-sided *p* < 0.01 threshold.

**Fig 12:**
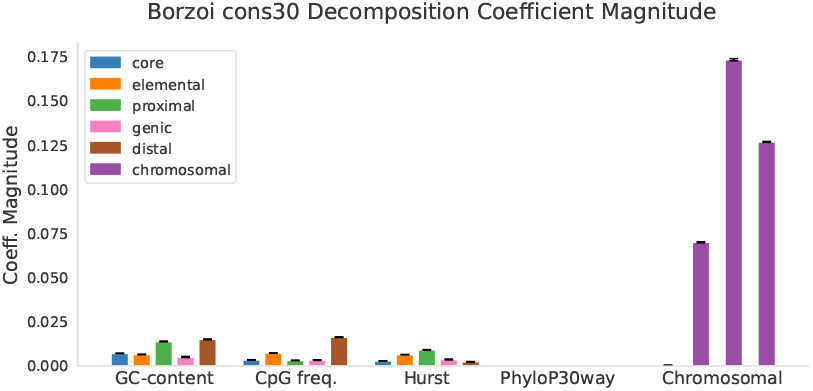
Coefficient magnitude (directionless) of the different features tested in the cons30 IVD of Enformer’s predictions. All features were standardized to mean zero and variance one before coefficient estimation. Error bars shown here are 3.259 times the test-set estimated standard error for each coefficient; this number corresponds to the Bonferroni-corrected two-sided *p* < 0.01 threshold. To prevent circularity issues, PhyloP30way features were not used here (features align vertically).

## Model comparative bin averaging resolution analyses

**Fig 13:**
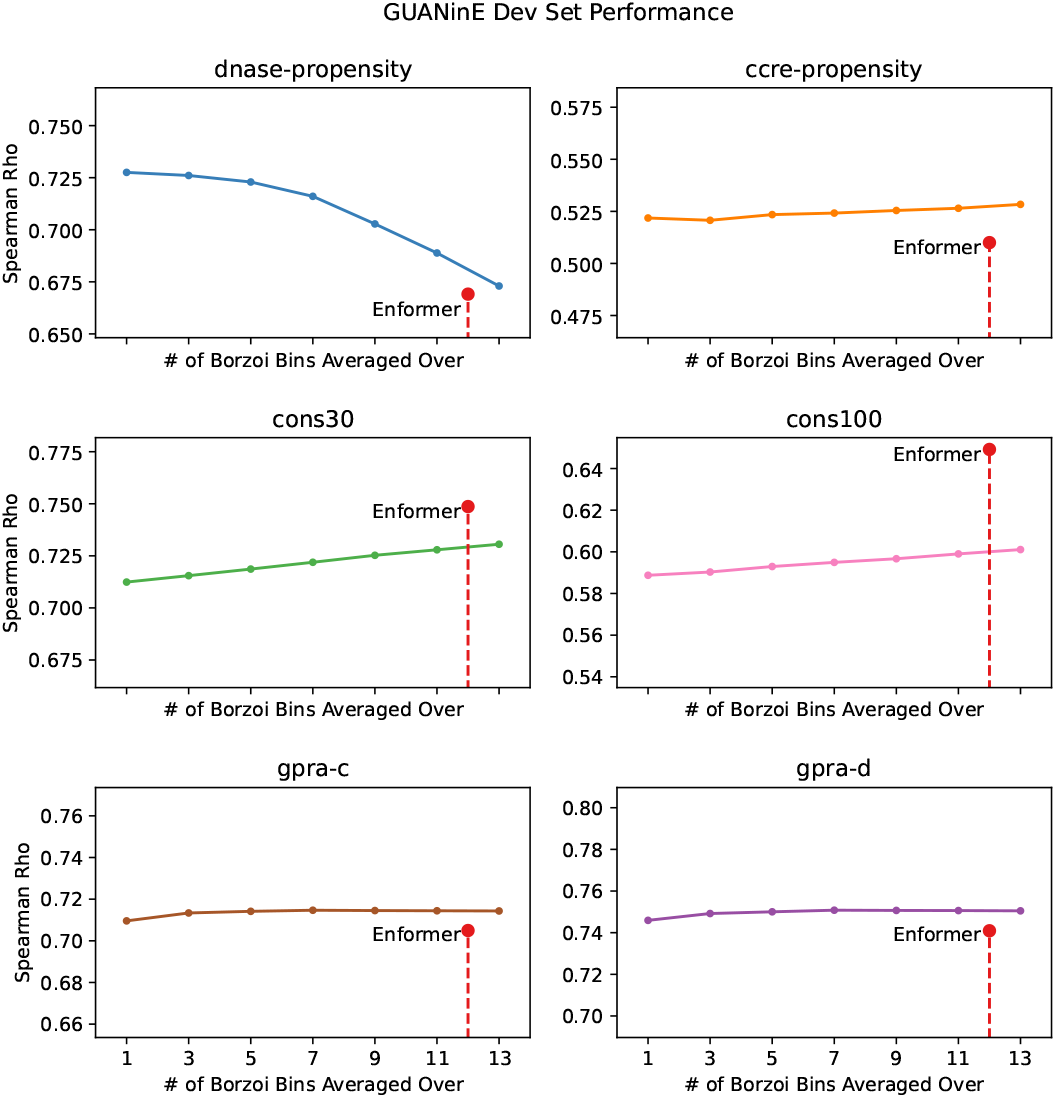
Development set performance of Borzoi on each `GUAN` in`E` task across multiple bin averaging size choices. Enformer’s dev set performance is shown relative to it’s 384 bp, which represents 12 bins.

## Footnotes

1 Though somewhat unwieldy, due to its input representation size, which is not well suited to modern deep learning hardware [42].

2 Cell-type-specific raw inputs are integrated to construct both cell-type-agnostic tasks. Note that Enformer’s 5,313-track and Borzoi’s 7,611-track training and validation sets include many of the individual input tracks.

3 The creators of the GPRA technique went to great lengths to design diverse sequences, and this increased variation shifts their distribution considerably from natural sequences [10, 54].

4 Compactness is a key concern, as the complete embeddings of Enformer’s training set total ∼ 1 T B and ∼ 1.4 *T B* for Enformer and Borzoi, corresponding to over one half and one quarter, respectively, of the size of the models’ outputs.

5 Both in terms of computational efficiency, given the cost of larger and larger models [39], and *statistical* efficiency, which governs how well a model fits, even in small sample sizes.

6 https://github.com/lucidrains/enformer-pytorch

7 https://huggingface.co/guanine

